# A Computational Framework to Optimize the Mechanical Behavior of Synthetic Vascular Grafts

**DOI:** 10.1101/2024.09.05.608688

**Authors:** David Jiang, Andrew J. Robinson, Abbey Nkansah, Jonathan Leung, Leopold Guo, Steve A. Maas, Jeffrey A. Weiss, Elizabeth M. Cosgriff-Hernandez, Lucas H. Timmins

**Author notes:** **Corresponding author** EnMed Tower, 1020 Holcombe Blvd., Houston, TX, 77030, U.S.A., *E-mail address* (L.H. Timmins).

## Abstract

The failure of synthetic small-diameter vascular grafts has been attributed to a mismatch in the compliance between the graft and native artery, driving mechanisms that promote thrombosis and neointimal hyperplasia. Additionally, the buckling of grafts results in large deformations that can lead to device failure. Although design features can be added to lessen the buckling potential, the addition is detrimental to decreasing compliance (e.g., reinforcing coil). Herein, we developed a novel finite element framework to inform vascular graft design by evaluating compliance and resistance to buckling. A batch-processing scheme iterated across the multi-dimensional design parameter space, which included three parameters: coil thickness, modulus, and spacing. Three types of finite element models were created in `FEBio` for each unique coil-reinforced graft parameter combination to simulate pressurization, axial buckling, and bent buckling, and results were analyzed to quantify compliance, buckling load, and kink radius, respectively, from each model. Importantly, model validation demonstrated that model predictions agree qualitatively and quantitatively with experimental observations. Subsequently, data for each design parameter combination were integrated into an optimization function for which a minimum value was sought. The optimization values identified various candidate graft designs with promising mechanical properties. Our investigation successfully demonstrated the model-directed framework identified vascular graft designs with optimal mechanical properties, which can potentially improve clinical outcomes by addressing device failure. In addition, the presented computational framework promotes model-directed device design for a broad range of biomaterial and regenerative medicine strategies.

## 1. INTRODUCTION

Nearly 400,000 coronary artery bypass grafting (CABG) procedures are performed annually in the U.S., making it the most common major surgical operation (Bachar & Manna, 2022). Although autografts are the standard of care for the procedure, they are unavailable in ∼10-20% of patients due to size mismatch, the need for multiple grafts, or peripheral vascular disease. Current synthetic vascular grafts such as expanded polytetrafluroethylene (ePTFE) or polyethylene terephthalate (Dacron™) have shown limited efficacy in small-diameter grafting (<6 mm) with a 40-50% reduction in patency after two years and 40% failing within five years (Salacinski, et al., 2001; Clowes, Gown, Hanson, & Reidy, 1985). The leading cause for the loss of patency of small diameter synthetic grafts is attributed to neointimal hyperplasia (NH) (Jeong, Yao, & Yim, 2021).

The development of NH has been correlated to a mismatch in the compliance, or radial stiffness, between the graft and native artery that can drive mechanisms promoting thrombosis and NH (Abbott, Megerman, Hasson, L’Italien, & Warnock, 1987). For instance, native arteries have compliance values ranging from 8 – 15 %/mmHg×10^−2^, permitting tissue dilation during the cardiac cycle (Abbott, Megerman, Hasson, L’Italien, & Warnock, 1987), whereas synthetic grafts have much lower compliance ranging compliance of 1.5-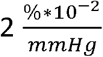 that results in reduced dilation (Greenwald & Berry, 2000). This compliance mismatch and corollary difference in vessel-graft dilation leads to a complex biomechanical environment characterized by disturbed blood flow patterns and wall stress gradients at the anastomoses (Szafron, Heng, Jack, Humphrey, & Marsden, 2024). A recent study demonstrated that disturbed flow patterns at the site of poor compliance matching between the graft and artery drive early markers of NH (Post, et al., 2019). Thus, compliance matching is a key biomechanical property for synthetic grafts to achieve long-term patency.

In addition to compliance matching, the mechanical stability of grafts under physiologic loads is essential to maintain their function. *Buckling*, defined as the loss of mechanical stability in a structure, results in large deformations of the material body that can lead to structural failure. Grafts can buckle under complex loads, resulting in well-characterized phenotypes (e.g., bent buckling, twist buckling, kinking) with important physiologic and clinical consequences (Han, Chesnutt, Garcia, Liu, & Wen, 2013). For a vascular graft, the loss of longitudinal stability by axial buckling can lead to a curved or tortuous shape, resulting in increased pressure loss and flow resistance that reduces distal perfusion (Wood, et al., 2006). Kinking, arising from bent buckling, can significantly restrict or completely occlude blood flow. Hence, it is crucial to design synthetic grafts to resist buckling and maintain functional performance.

Important to device design is recognizing that optimizing one mechanical characteristic could negatively impact another (i.e., mechanical attributes could be in counter-balance). For example, studies show that vascular grafts with high compliance can buckle after implantation (Zizhou, Wang, & Houshyar, 2022). ePTFE or Dacron™ grafts contain unique design features that prevent kinking but at the consequence of low compliance (Heng, et al., 2023). Recognizing the compliance matching capability of hydrogel-electrospun vascular grafts (Post, et al., 2019), but understanding their potential to buckle, it has been proposed to add a reinforcing coil around the construct to increase the buckling resistance (Li X., 2018). However, this addition could negatively impact the compliance of the graft. Based on geometric and material property tunability, the number of design parameter combinations to balance the opposing properties in a coil-reinforced graft is nearly limitless and prohibitively time-consuming to manufacture via a trial-and-error approach.

Herein, we develop a novel finite element (FE) modeling framework to inform vascular graft design by iterating across the multi-dimensional design parameter space, simulating the relevant biomechanical loading conditions, and evaluating compliance and resistance to buckling. Fabricated synthetic vascular grafts with coil supports are utilized to compare experimental and simulation results to validate the models. Integrating the framework into an optimization scheme provides an approach to identify candidate graft designs that reconcile three competing solid mechanical concerns – compliance, axial buckling, and bent buckling.

## 2. MATERIAL AND METHODS

### 2.1 Parametric Design Space

Based on prior evidence of competing mechanical properties between buckling resistance and compliance for a coil supported graft, coil design parameters were investigated to balance the two properties (Figure 1a). Coil designs were uniquely defined using the following three parameters: coil thickness (*δ*) ranged from 0.1 - 0.5 mm (5 equally-spaced values), coil modulus (*μ*) of two different materials (Bionate® 80A and 55D), and coil spacing (*p*) ranged from 1 - 10 mm (10 equally-spaced values). Across these design input parameters, 100 unique coil-reinforced graft combinations were examined (Figure 1b).

**Figure 1.**
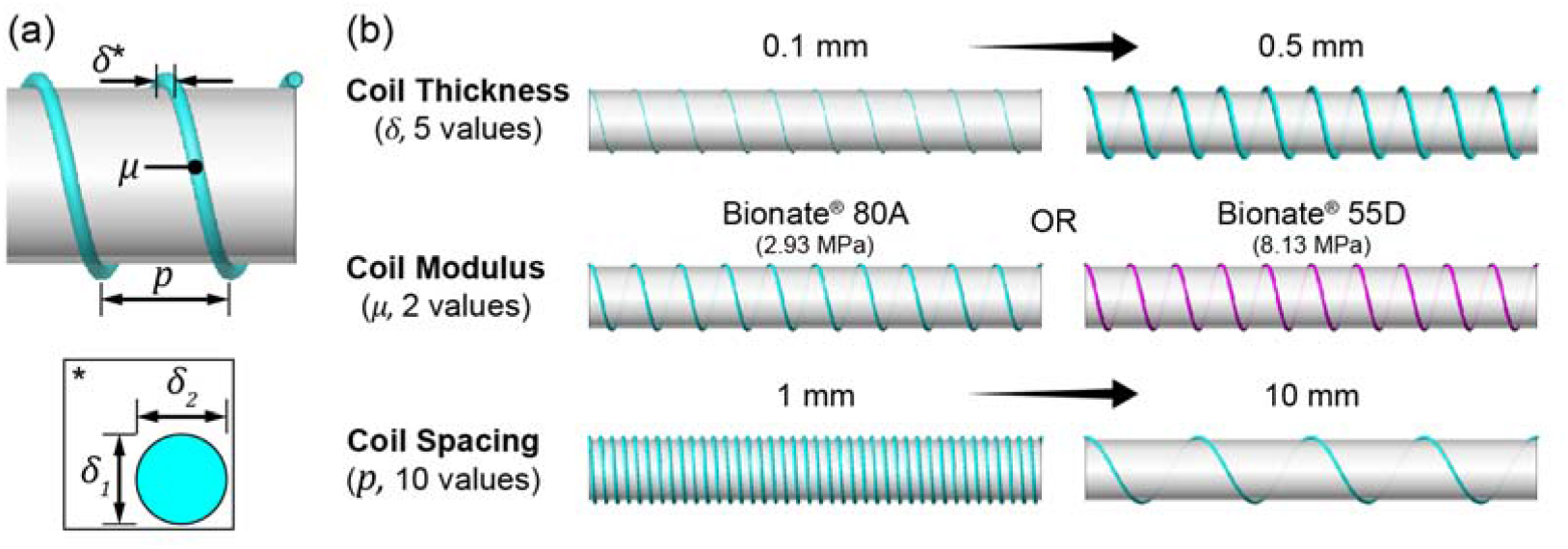
Coil-reinforced graft design parameters. (a) Reinforcing coil (right-handed, cylindrical helix) showing the three parameters of interest: *δ* is the thickness of the coil, *μ* is the shear modulus of the coil material, and *p* is the pitch or height of one complete helix turn. (b) Renderings of the coil-reinforced grafts by varying the three design parameters. [1.5-column]

### 2.2 FE Model Development

All FE models were developed in the open-source FE software `FEBio` (“Finite Element for Biomechanics”) and `FEBio Studio`^2^ (Maas, Ellis, Ateshian, & Weiss, 2012; Maas, Ateshian, & Weiss, 2017). To accelerate model development, a batch-processing scheme was developed in `MATLAB` (MathWorks, Natick, MA) to sweep the multi-dimensional parameter space. The automated subroutine created and discretized the model geometries, applied model definitions (i.e., boundary conditions, loads, contact), and generated `FEBio` input files.

#### 2.2.1 Geometry & Mesh

The automated scheme utilized the open-source `GIBBON` toolbox^3^ to construct and discretize the coil-reinforced graft geometry (Moerman, 2018). The grafts were modeled as an axisymmetric cylinder with a reinforcing coil wrapped helically around the outer graft surface (Figure 2). Beyond the three coil parameters described in section 2.1, additional properties of the coil-reinforced graft model are defined in Table 1.

**Table 1.** Coil-reinforced graft model characteristics and dimensional values.

| Model Parameter | Description | Value(s) |
| --- | --- | --- |
| $D_i$ | Graft inner diameter | 5.0 mm |
| $D_o$ | Graft outer diameter | 5.5 mm |
| $L$ | Graft length | 40 mm |
| $\delta$ | Coil thickness | 0.1 – 0.5 mm |
| $\mu$ | Coil shear modulus | 2.93, 8.13 MPa |
| $p$ | Coil pitch | 1 – 10 mm |
| $\theta$ | Coil start angle | 0, 90, 180, 270° |
| $\alpha_0$ | Imperfection angle | 0° (i.e., none) |

**Figure 2.**
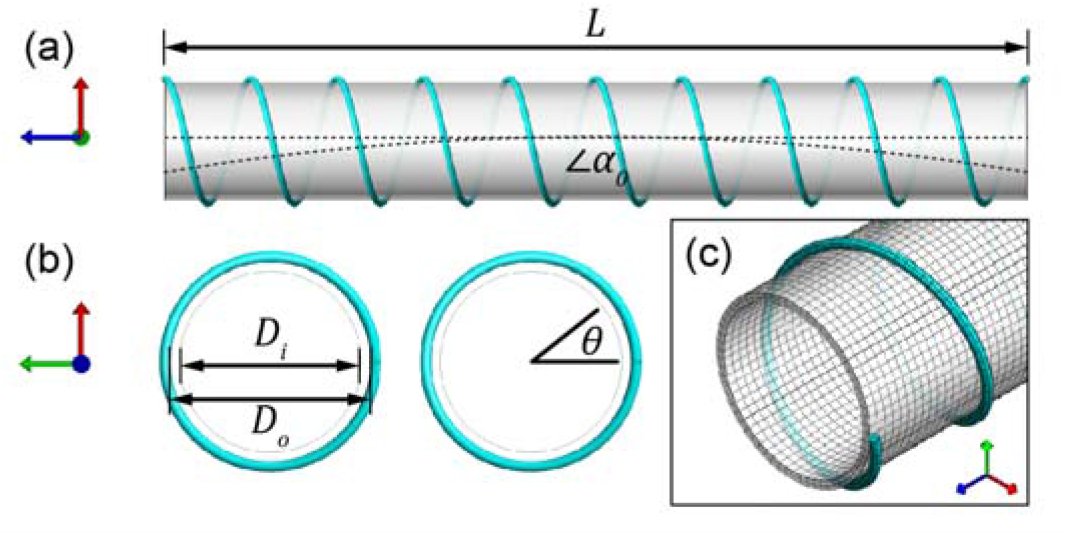
Geometric dimensions of the coil-reinforced graft model. (a) Side view of the model with associated dimensions for graft length (*L*) and imperfection angle (*α*_0_, drastically increased for visual clarity) (b) Top view associated of the model with associated dimensions for the graft inner and outer diameters (*Di, D*_*o*_), and *θ* defines where the coil turn starts. (c) Isometric view of the meshed graft and coil parts, composed of hexahedral and pentahedral elements, respectively. [single-column]

The creation of the model is described as follows: First, a sketch was created of a regular polygon with the diameter, *Di*. Next, the 2D sketch was extruded to obtain surface patch data (i.e., quadrilateral surface mesh),generating a CAD-like model geometry of a tube with length *L*. The surface was then thickened to the outer diameter, *D*_*o*_, and solid hexahedral elements were generated from the patch data. Subsequently, a ketch of the coil cross-section was created with a thickness of *δ*, and a guide curve of a helix was defined with a pitch between turns of length *p* and a start angle *θ*. The sketch was swept along the guide curve while ensuring no twisting occurred. The components were merged to create a quadrilateral surface mesh of the coil and discretized into solid pentahedral elements following a right-handed node numbering convention. After obtaining the nodal coordinates of the FE mesh, an in-plane imperfection was specified to displace the undeformed geometry with an initial bend, expressed by a factor *α*0, along the central axis to facilitate the first buckling mode (Sanyal & Han, 2015).

The graft and helical coil geometries were discretized with tri-linear 8-node hexahedral elements and 6-node pentahedral elements, respectively, and utilized a three-field element formulation (Simo & Taylor, 1991). The graft domain consisted of 18,816 structured elements, and the coil domain comprised of 2,392 – 22,368 structured elements, depending on the coil pitch (Figure 2c). A mesh convergence study was performed based on the criteria of <3% change in evaluated metrics (i.e., compliance, buckling load, and kink radius)^4^. Finally, surface features (e.g., top face, inner face, etc.) were automatically detected from the grouping of geometric patch data based on a maximum dihedral angle to aid the application of boundary conditions, loads, and contact.

#### 2.2.2 Constitutive Model

The graft and coil material properties were derived from mechanical testing and Bionate® PCU product sheets (DSM Biomedical Inc). Each was modeled in `FEBio` as an *uncoupled neo-Hookean* hyperelastic material with nearly incompressible behavior, which uncouples the neo-Hookean constitutive model into a deviatoric and volumetric behavior, given by the strain-energy function,

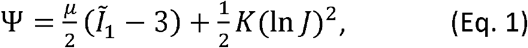

where *μ*is the shear modulus material coefficient, Ĩ_1_is the first invariant of the deviatoric right Cauchy-Green deformation tensor, *K* is a bulk modulus-like penalty parameter, and *J* is the determinant of the deformation gradient tensor. The shear modulus for the graft mesh was 671 kPa, and the shear moduli for the two coil materials are shown in Figure 1b. Limited compressibility (i.e., ν≈ 0.45) for all materials was incorporated using the bulk modulus, which was defined as 10× the shear modulus. The densities for the graft and coil were 1090 kg/m^3^ and 1200 kg/m^3^, respectively.

#### 2.2.3 Boundary Conditions & Loads

Three types of FE models were developed for each unique coil-reinforced graft to simulate (a) pressurization, (b) axial buckling, and (c) bent buckling (Figure 3). For all models, both graft ends were fixed to only undergo rigid body motion (i.e., affine translation and rotation) during the entire analysis. Specifically, boundary conditions were applied to all nodes within a reference point coupled to 3 mm sections at each end of the graft.

**Figure 3.**
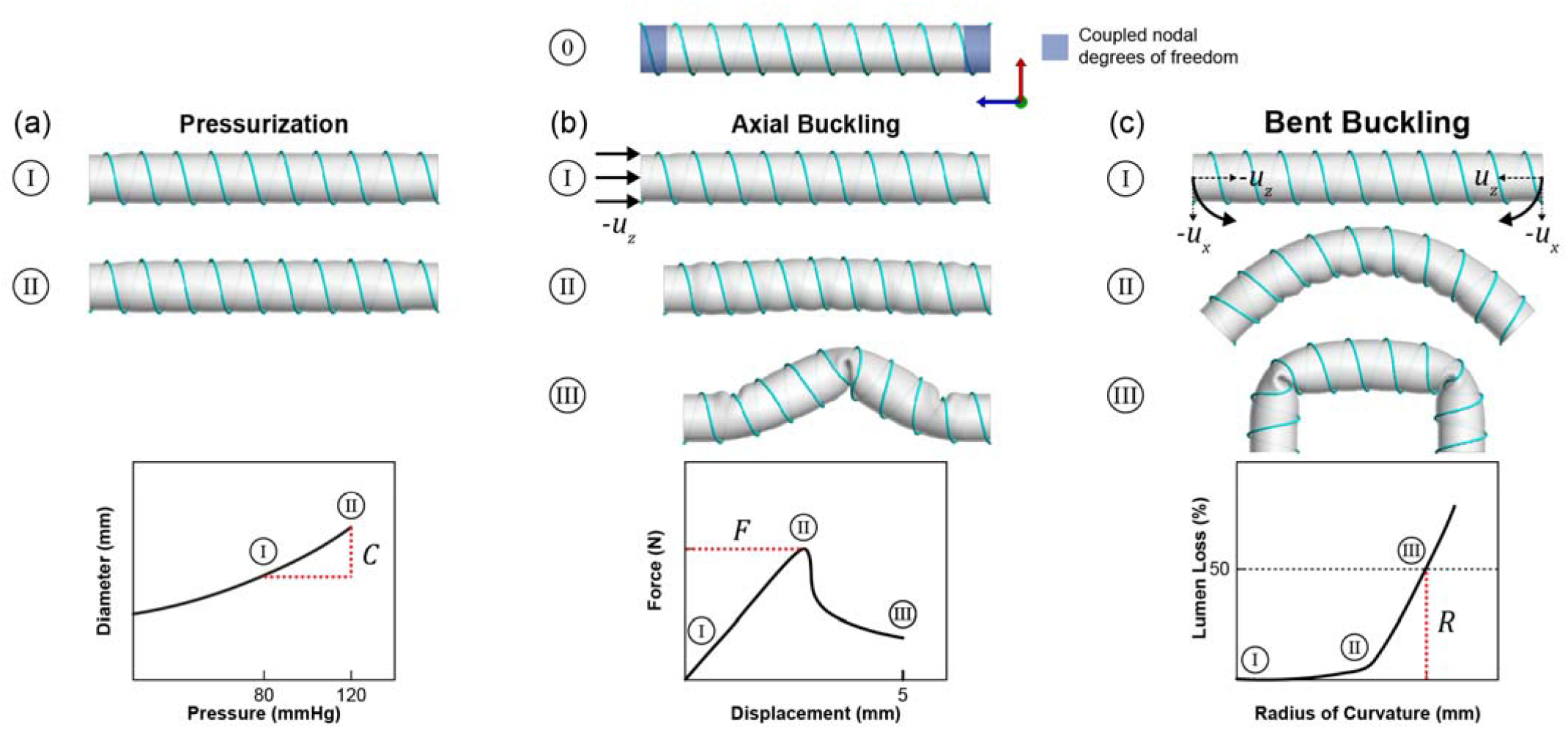
Illustrations of coil-reinforced graft deformation procedure. In the initialization phase (0), all nodes highlighted in the blue regions are coupled together and remain active for all subsequent steps. (a) Pressurization up to 80 (I) and 120 mmHg (II) caused a change in the diameter of the graft. (b) Axial buckling due to pressurization of 100 mmHg (I) and subsequent displacement to compress the graft (II) prompted a lateral deflection (i.e., loss of stability) of the structure (III). (c) Bent buckling due to pressurization of 100 mmHg (I) and subsequent displacements to bend the graft (II) formed a local kink or discontinuity in the structure (III). [2-column]

1. To perform pressurization, a *zero-displacement* boundary condition fixed both graft ends in all three degrees of freedom (DOF) to prevent rigid body modes (i.e., *u*_*x*_ = *u*_*y*_ = *u*_*z*_ = 0). A *pressure* load (*P*) was applied to the graft inner surface and inflated to physiologic pressures of 80 and 120 mmHg (Figure 3a).
2. To perform axial buckling, the graft ends were fixed in all DOFs. Deformation of the graft occurred in two stages, a pressurization regime (*T*_1_) and a displacement regime (*T*_2_). The graft inner surface was pressurized and held constant at 100 mmHg. Subsequently, a *prescribed-displacement* boundary condition (−u_z_) was applied normal to one end of the graft, compressing the graft longitudinally by 5 mm (Figure 3b).
3. To perform bent buckling, the graft ends were fixed in all DOFs. As in (b), deformation of the graft occurred in two stages, a pressurization regime (*T*_1_) and a displacement regime (*T*_2_) The graft inner surface was pressurized to 100 mmHg. Next, *prescribed-displacement* boundary conditions were applied equally and in opposite rotations to each end of the graft, displacing the ends downwards (−u_x_) and inwards (*u*_z_) to generate an effective bending moment (Figure 3c). Essentially, these boundary conditions applied the kinematics of pure bending to decrease the radius of curvature of the central graft axis to 4 mm. Strategies to simulate bent buckling were modified from methods for kink deformation described in (Mckenna & Vaughan, 2021).

For models with a coil, a *tied-elastic* contact interface defined the interaction between the graft outer-surface and the coil surface. This contact formulation connected (i.e., bonded) the two non-conforming meshes by enforcing continuity of displacement across the interface (Laursen & Simo, 1993). The *sliding-elastic* contact formulation was prescribed at multiple interfaces across the graft and coil components to prevent self-penetration of the elastic bodies during deformation (Zimmerman & Ateshian, 2018). This boundary condition was applied across the following: (i) graft outer-surface to graft outer-surface, (ii) graft outer-surface to coil surface, and (iii) coil surface to coil surface. The non-penetrating constraints were enforced via an auto-penalty method, updated at each iteration to account for the deformation. To alleviate convergence problems due to multiple folds on the contact surfaces, a *search radius* of *δ*/2 was defined so that the algorithm ignored contact pairs outside the dimensional search radius.

#### 2.2.4 Simulation Settings

Each coil-reinforced graft configuration was simulated using a combination of the implicit and explicit solid solvers in `FEBio`. The implicit solver simulated the graft’s pressurization regime (*T*_1_= 0.1 s), efficiently Each coil-reinforced graft configuration was simulated using a combination of the implicit and explicit solving the low-strain, quasi-static deformation. The implicit solver control parameters were as follows: time steps = 10, time step size = 0.01, displacement tolerance = 0.001, energy tolerance = 0.01, linear search tolerance = 0.9, update method: full Newton, global stiffness matrix: unsymmetric, linear solver: Pardiso. The explicit solver was employed for the axial buckling and bent buckling models, which was preferred for the complex geometric, material, and contact non-linearities caused by the dynamic instabilities of the graft buckling^5^. The computational cost for the explicit model was reduced via load-rate scaling to artificially increase the rate of applied boundary conditions over a shorter period (*T*_2_ = 0.1 s) since the FE models do not possess rate-dependent materials. The load rate was limited to less than 1% of the wave speed of the graft. As explicit-time integration schemes are only conditionally stable, a time step size of 2e-07 s was identified for 2015). In addition, a mass damping scheme was employed with a damping coefficient (*C*= 300 Ns/m) to the solution to remain valid, resulting in a total of 5e+05 timesteps (Askes, Rodriguez-Ferran, & Hetherington, 2015). In addition, a mass damping scheme was employed with a damping coefficient (*c* = 300 Ns/m) to suppress the vibrational modes and to obtain the equilibrated solution. To ensure a quasi-static solution, the ratio of kinetic energy to internal strain energy did not exceed 5% throughout the analysis^6^.

FE simulations were solved using a high-performance computing (HPC) system, containing 16 Intel^®^ Xeon^®^ Platinum 8360H processors (384 cores) @ 3.00-4.20 GHz, and 3TB of DDR4-3200 RAM (Scientific Computing and Imaging Institute, University of Utah). Multiple simulations were solved simultaneously with a single core designated per job. Following model convergence, output files were automatically post-processed for analysis in `MATLAB` and visualized in `FEBio Studio`.

### 2.3 Evaluation Methods

#### 2.3.1 In Silico Metrics

The FE simulation results were analyzed for three biomechanical metrics: (a) compliance due to pressurization, (b) buckling load due to axial buckling, and (c) kink radius due to bent buckling. The procedures described in ISO 7198:2016 guided the analysis methods and subsequent experimental testing, further described in section 2.4.3.

a Compliance (*C*, units of %/mmHg×10^−2^) was quantified by measuring the change in diameter due to a change in pressure (Figure 3a). That is,

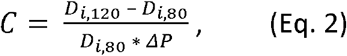

where *D*_*i*,80_ and *D*_*i*,l20_ are the inner graft diameters at 80 mmHg and 120 mmHg, respectively, and Δ*P* is the change in pressure. The inner diameters at 25%, 50%, and 75% along the graft length were extracted from deformed nodal positions to calculate mean graft compliance. Target values of graft compliance in better agreement with arterial tissue indicated greater compliance matching properties.

b The buckling load (*F*, units of N) was quantified as the maximum compressive load the graft supported under an applied axial displacement (Figure 3b). Nodal values of the reaction force were extracted at the location of applied displacement and summed. Higher values of buckling load indicated greater resistance to axial buckling.
c The kink radius (*R*, units of mm) was quantified as the inner radius of curvature required to kink the vascular graft (Figure 3c). A kink was defined as a >50% greater reduction in inner diameter during the deformation (International Organization for Standardization, 2016). Nodal positions were extracted from the graft inner surface and in-plane to the direction of applied bending to determine the presence of a kink. Lower values of kink radius indicated greater resistance to bent buckling.

#### 2.3.2 Optimization Function

Data from the FE models across the coil design parameter space were integrated into an optimization function. Across the 100 models, the values for compliance, buckling load, and kink radius were standardized.

Since a compliance matching graft is optimal, graft compliance values (*C*_*graft*_) were normalized against the reported value for coronary tissue (*C*_artery_, 8 %/mmHg×10^−2^, (Abbott, Megerman, Hasson, L’Italien, & Warnock, 1987)). As a high buckling load is preferred, buckling loads (*F*_*graft*_) were standardized against the minimal buckling load across the models. As a low kink radius is favorable, kink radius values (*R*_*graft*_) were standardized against the maximum kink radius across the models. A minimal value was sought for the, optimization function (φ), which was defined as

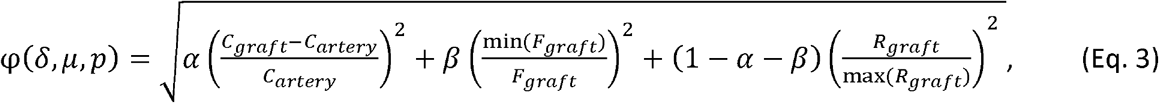

where *δ, μ*, and *p* are the input parameters – coil thickness, coil modulus, and coil pitch, respectively. The weighting coefficients *α* and *β* represent the relative importance of the compliance, buckling load, and kink radius. Various *α* and *β* values were investigated.

### 2.4 Graft Fabrication and Experimental Testing

A validation study was performed to promote FE model confidence. Three graft geometries were fabricated to allow comparison between experimental and FE-derived results. All reagents were obtained and used as received from Sigma-Aldrich (Milwaukee, WI) or VWR (Radnor, PA) unless stated otherwise.

#### 2.4.1 Electrospinning

Vascular grafts were fabricated via electrospinning as described previously (Post, et al., 2019). A 14wt% solution of Bionate® 80A (DSM Biomedical) in a solvent mixture of 70:30 dimethylacetamide:tetrahydrofuran was passed through a 20-gauge blunt needle at a flow rate of 0.5 ml/hr and charged to +15.3-22.5 kV. A 5 mm rod was dip coated in a solution of 5wt% polyethylene glycol (PEG; 35 kDa) in dichloromethane to facilitate graft removal following spinning and rotated at 500 RPM. The mandrel was placed 50 cm from the needle and charged to –2.5 kV. Relative humidity and temperature were 40-48% and 23.2-25.4 °C, respectively. Electrospun control grafts were spun to a thickness of ∼250 µm. Coil-reinforced grafts were fabricated in three layers with an initial electrospun layer of ∼120-160 µm. A polymer coil was then applied using a custom printer to deposit an 18 wt% solution of Bionate® 55D in 1,1,1,3,3,3-Hexafluoro-2-propanol. Finally, a second electrospun layer was applied to the graft for a total mesh thickness of ∼250 µm. Grafts were removed by placing the mandrel in water for 30 minutes to dissolve the PEG layer and trimmed to ∼4 cm. To improve layer integrity, the grafts underwent solvent vapor welding by exposure to tetrahydrofuran solvent vapor for two hours prior to testing.

#### 2.4.2 Hydrogel Coating

A polyether urethane diacrylamide (PEUDAm, 20 kDa) macromer was synthesized using established protocols (Wancura, et al., 2023). Electrospun polyurethane grafts were coated with the PEUDAm hydrogel using diffusion-mediated redox crosslinking, as described previously (Wancura, Talanker, Toubbeh, Bryan, & Cosgriff-Hernandez, 2020). Briefly, electrospun grafts first underwent an ethanol-water wetting ladder (70 vol%, 50 vol%, 30 vol%, and 0 vol% ethanol in water, with each concentration applied for 15 minutes). Grafts were then submerged in iron gluconate dihydrate solutions (IG, 3 wt/vol% [Fe^2+^], as determined with the Ferrozine Assay, (Jeitner, 2014)) for 15 minutes, briefly dipped in methanol, and air dried for one minute. The IG-coated substrates were then immersed in aqueous solutions of 10 wt/vol% PEUDAm with ammonium persulfate (APS, 0.05 wt/vol%) and 2 wt/vol% Irgacure 2959 for 20 seconds. Grafts were removed and placed on a UV plate (Intelligent Ray Shuttered UV Flood Light, Integrated Dispensing Solutions, Inc., 365 nm, 4 mW/cm^2^) for an additional photocure period of 12 minutes. Grafts were dried overnight and soaked for 5 hours in DI water, with water exchanges performed after 10 minutes, 1 hour, and 3 hours, followed by redrying of the grafts. Finally, hydrogel-coated grafts were immersed in a solution of 20 wt/vol% N-acryloyl glycinamide (NAGA, BLD Pharma), 0.1 mol% bisacrylamide (bisAAm), and 2 wt/vol% Irgacure 2959 overnight protected from light at 4 °C. The grafts were then placed on a 4 mm borosilicate glass rod and placed on a UV plate (Intelligent Ray Shuttered UV Flood Light, Integrated Dispensing Solutions, Inc., 365 nm, 4 mW/cm^2^) for 12 minutes to initiate photopolymerization of NAGA as a second interpenetrating network. The final composite grafts were dried overnight and swelled to equilibrium in DI water prior to testing.

#### 2.4.3 Graft Experimental Testing

A control group of grafts with no support coil was fabricated (unsupported group, *n*= 3), as well as two groups of coil-reinforced grafts: supported group 1 (*δ* = 0.15 mm, μ = 55D, *p* = 4.5 mm, *n*= 4) and supported group 2 (*δ* = 0.35 mm, μ = 55D, *p*= 2.5 mm, *n*= 2). The experimental testing methods were based on ISO 7198:2016, which specified requirements for evaluating vascular prostheses based on current medical knowledge (International Organization for Standardization, 2016).

a. To determine compliance, grafts were pressurized up to 120 mmHg at a rate of 2 ml/min using water at room temperature. Graft outer diameter at 8 positions was measured using a laser micrometer at pressures of 80 mmHg and 120 mmHg (Z-Mike 1210 Laser Micrometer). Compliance was calculated as defined in Eq. 2.
b. The buckling load was measured using a custom, computer-controlled experimental test platform. Utilizing a syringe pump, multi-axis motional controllers, load cell, and CCD camera, mounted grafts were pressurized to 100 mmHg and compressed 10 mm to induce buckling. Displacement control and force measurements (*n*= 3 per graft) were controlled with `LABVIEW`.
c. Kink radius tests were performed on 3D printed cylindrical mandrels, with diameters ranging from 41 to 5 mm in increments of 3 mm (International Organization for Standardization, 2016; Wu D. J., et al., 2020). Grafts were pressurized to 100 mmHg and bent to the curvature of each mandrel while ensuring the mandrel did not support the graft. The mandrel radius was decreased until the graft was kinked, as defined in Section 2.3.1. Measurements (*n* = 3 per graft) were made and analyzed in `ImageJ`.

FE geometries were created, as described in Section 2.2, with the same design parameters as those fabricated (*D*_*i*_ = 4.5 mm, *D*_*o*_ = 5.0 mm, *L*= 40 mm). To account for variation in coil orientation during experimental testing, FE models were created where the circumferential position of the coil origin varied (i.e., models were created where *θ* in Table 1 and Figure 2 was defined as 0°, 90°, 180°, and 270°). The shear modulus for the graft mesh was 610 kPa. Recognizing that imperfections exist within the manufactured grafts (e.g., geometric tolerances, fabrication defects) that could skew validation results, an initial imperfection was incorporated into the model (i.e., where *α*_*o*_ in Table 1 and Figure 2 was defined as 5°). Boundary conditions were applied to all nodes within a reference point coupled to 5 mm sections at each end of the graft. FE simulations were solved, and model output metrics were analyzed as described in Section 2.3.1.

### 2.5 Statistical Analysis

FE model and experimental data analyses were performed in `MATLAB`. Descriptive statistics are reported as mean ± standard deviation. Significance was determined using a one-sample student’s *t*-test, and statistical significance was defined as a *p*-value < 0.05. In addition, Cohen’s *d* was calculated to measure the effect size, an indication of the strength in a statistical relationship, and values were classified as trivial (0 ≤ *d* < 0.2), small (0.2 ≤ *d* < 0.5), medium (0.5 ≤ *d* < 0.8), or large (*d* ≥ 0.8). To understand the influence of changes in design parameters across the wide ranges, a sensitivity analysis was performed to characterize and understand the influence of graft design parameters(*δ, μ,p*) on the predicted biomechanical response metrics (*C,F,R*)Sensitivity (*S*) was defined as,

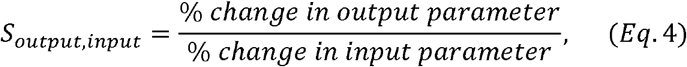

where the numerator is the mean percent change in the model output and the denominator is the mean percent change in the model input parameter.

## 3 RESULTS

### 3.1 FE Model Validation

Examination of three vascular graft groups demonstrated qualitative agreement in the deformed geometries between the FE-predicted and experimental data, as well as quantitative agreement in compliance, buckling load, and kink radius (Figure 4). Specific to each loading condition,

**Figure 4.**
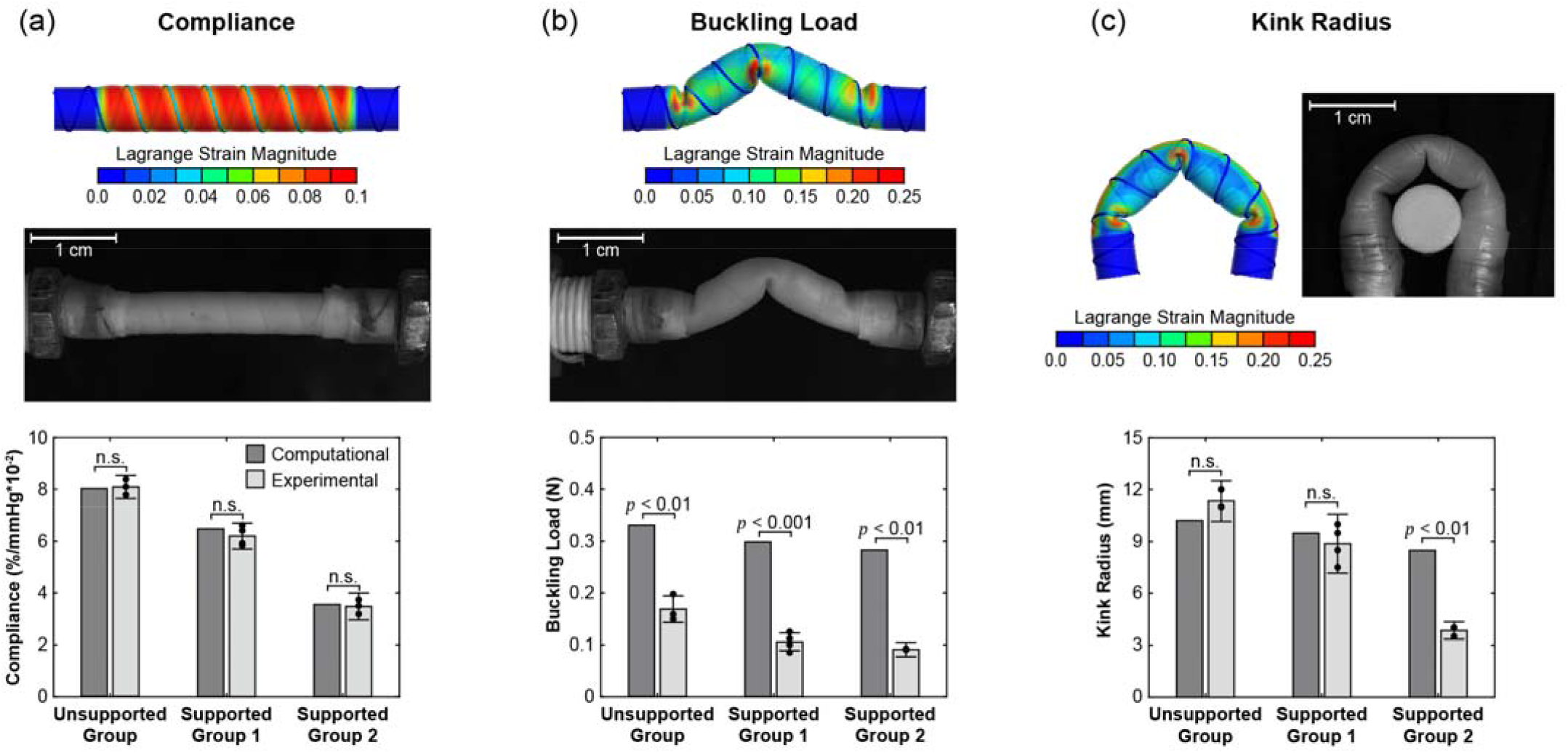
Comparison of computational FE model prediction and experimental measurements of three graft groups: unsupported group (no coil), supported group 1 (*δ*= 0.15 mm, _*μ* = 55D, *p* = 4.5 mm), and supported group 2 (*δ* = 0.35 mm, *μ* = 55D, *p* = 2.5 mm). Deformations of the graft were simulated and tested to analyze (a) compliance due to pressurization, (b) buckling load due to axial buckling, and (c) kink radius due to bent buckling. [double-column].

a. The FE results from the compliance models highlighted the presence of lower Lagrange strains near the coils, as compared to the regions that did not have coil reinforcement (i.e., the coil restricted graft deformation). The experimental testing confirmed these findings. Modeling data demonstrated that the addition of the coil decreased compliances from 8.0 %/mmHg×10^−2^ (unsupported group) to 6.5 %/mmHg×10^−2^ (supported group 1) and 3.6 %/mmHg×10^−2^ (supported group 2). Likewise, experimental data demonstrated the addition of the reinforcing coil reduced the compliance from 8.1 ± 0.5 %/mmHg×10^−2^ (unsupported group) to 6.2 ± 0.5 %/mmHg×10^−2^ (supported group 1) and 3.5 ± 0.5 %/mmHg×10^−2^ (supported group 2). No significant differences were determined between the FE models and experiments in all three groups. Importantly, effect sizes for the compliance across the groups were 0.15 (unsupported group), 0.57 (supported group 1), and 0.15 (supported group 2), indicating small to medium differences. The relative error between the FE results and experiments was 0.1 ± 0.2 %/mmHg×10^−2^.
b. In the axial buckling models, the FE results showed three kinks formed along the graft, as indicated by high Lagrange strains in the middle of the graft and ends. The deformation in the experiments supported the predicted FE model behavior, as the kinks formed in approximately the same locations. FE model predicted buckling loads exhibited minor differences across the three groups with values of 0.33 N (unsupported group), 0.30 N (supported group 1), and 0.28 N (supported group 2), highlighting that the addition of the coil decreased the buckling load. Experimental data also showed that the addition of the coil decreased the buckling load with minor differences with values of 0.17 ± 0.03 N (unsupported group), 0.11 ± 0.02 N (supported group 1), and 0.09 ± 0.01 N (supported group 2). Significant difference was observed in all three groups. The effect sizes across the three groups were 8.45 (unsupported group), 10.25 (supported group 1), and 11.42 (supported group 2), indicating large differences. However, the relative error was consistent between the FE predictions and experiments - the FE models over-predicted buckling load by 0.18 ± 0.02 N.
c. The FE results of the bent buckling model showed three kinks formed along the graft that were also observed experimentally. Moreover, the FE models captured the experimentally observed trend of reduced kink radius with the addition of a supporting coil. The model-predict kink radius decreased from 10.2 mm (unsupported group) to 9.5 (supported group 1) and 8.5 mm (supported group 2) when a coil was added. Experimental measurements of kink radius showed a reduction from 11.3 ± 1.2 mm (unsupported grafts) to 8.9 ± 1.7 mm (supported group 1) and 3.8 ± 0.5 mm (supported group 2). No significant differences were determined in the unsupported group and supported group 1. The effect sizes were 1.05 (unsupported group), 0.36 (supported group 1), and 5.90 (supported group 2), indicating small to large differences.

### 3.2 *In Silico* Evaluation of Biomechanical Metrics Across Graft Design Parameter Space

The compliance, buckling load, and kink radius varied non-linearly across the parameter design space (Fig. 5), and sensitivity analysis further demonstrated dependencies between model output metrics and design parameters (Table 2). Specific to each loading condition,

**Table 2.** Mean sensitivity values of output computational metrics with respect to input design parameters.

|  | Compliance | Buckling Load | Kink Radius |
| --- | --- | --- | --- |
| Coil thickness | $S_{C,\delta} = 0.23$ | $S_{F,\delta} = 0.11$ | $S_{R,\delta} = 0.13$ |
| Coil modulus | $S_{C,\mu} = 0.07$ | $S_{F,\mu} = 0.04$ | $S_{R,\mu} = 0.03$ |
| Coil pitch | $S_{C,p} = 0.27$ | $S_{F,p} = 0.61$ | $S_{R,p} = 0.39$ |

**Figure 5.**
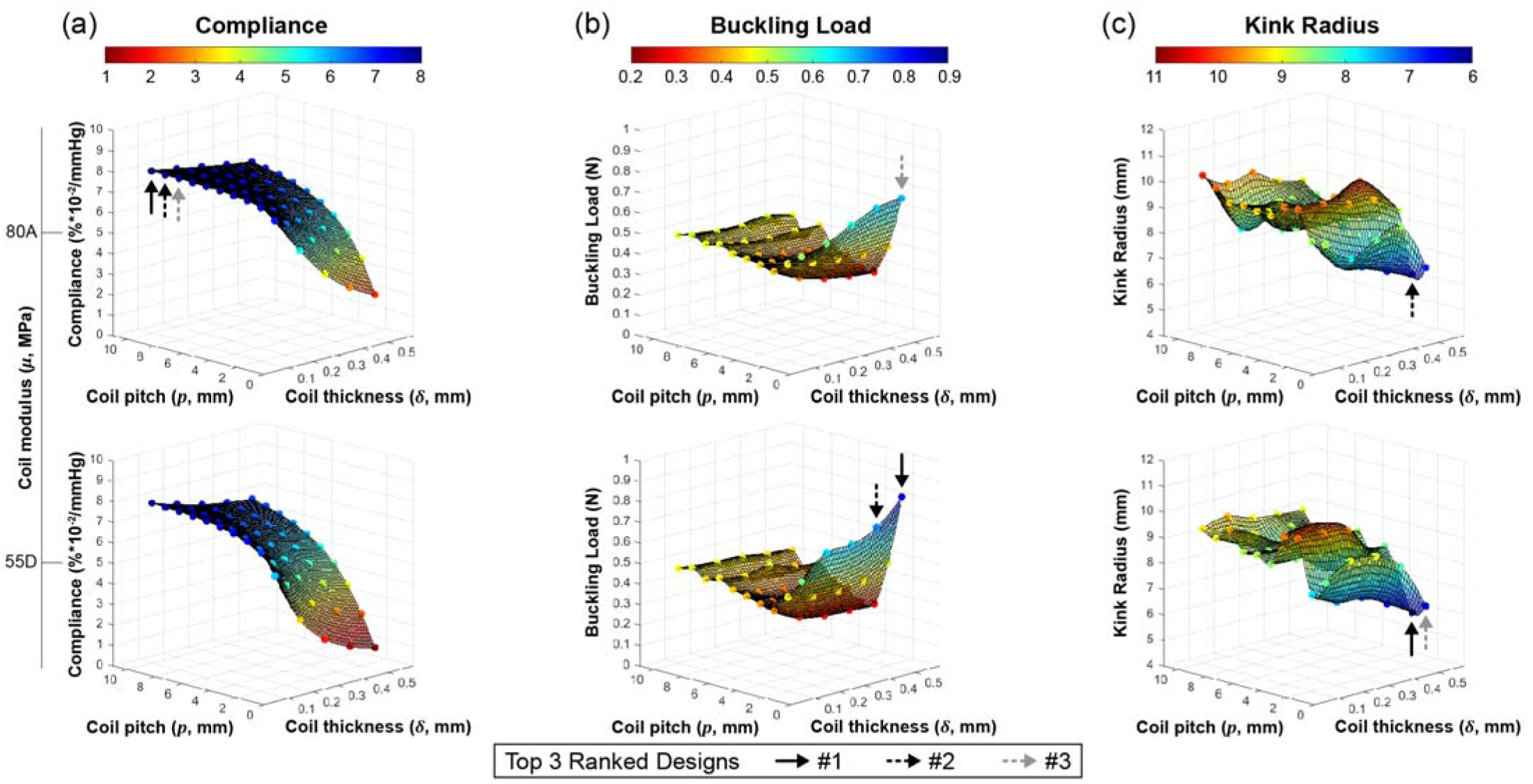
Design space evaluations for the 100 unique coil-reinforced graft combinations. Data were interpolated within the design space to evaluate (a) compliance, (b) buckling load, and (c) kink radius. Each output metric was plotted against the – coil thickness (*δ*) and coil pitch (*p*). The top and bottom rows separately represent the two coil moduli (*μ*), Bionate ® 55D and 80A, respectively. Optimal mechanical properties are identified by the arrows representing the top 3 ranked designs for each output metric.

a. The graft compliance displayed a wide range of values from 1.1 to 7.9 %/mmHg×10^−2^ across the 100 evaluated models (Figure 5a). Values decreased unidirectionally across the parameter space, with increasing coil thickness and decreasing coil pitch. Notably, the variation in compliance increased as coil pitch decreased. At a coil pitch of 10 mm, the compliance was 7.4 ± 0.4 %/mmHg×10^−2^. At a coil pitch of 1 mm, the compliance was 3.5 ± 2.0 %/mmHg×10^−2^. Compliance values prominently for coil pitch values of *δ*< 4 mm. Sensitivity analysis demonstrated that the compliance values were slightly more dependent on coil thickness than pitch, with minimal changes in compliance across coil moduli (Table 2). Across all pressurization models, the mean solve time was ∼3 min, and the longest model took 4 min.
b. The buckling loads across the parameter space ranged from 0.28 to 0.84 N (Figure 5b). While buckling loads had minor variations from coil pitches of 3 to 10 mm, the buckling load increased sharply at coil pitches of 2 and 1 mm. For instance, at a coil pitch of 3 mm, the mean buckling load was 0.32 ± 0.05 N, whereas, at coil pitches of 2 and 1 mm, the mean buckling load was 0.45 ± 0.02 N and 0.65 ± 0.09 N, respectively. Sensitivity analysis supported these observations, showing a high dependence on coil pitch, followed by coil thickness and modulus (Table 2). Across all axial buckling models, the solve time was 04:15:26 ± 01:02:28 (days:hours:minutes), and the longest model took ∼8 days to complete.
c. The kink radius varied from approximately 6 to 10 mm in the evaluated models (Figure 5c). Across the design space, the kink radius fluctuated unpredictably between coil pitch values of 4 to 10 mm, in addition to oscillations across the coil thickness. However, at a coil pitch of 3 mm, the kink radius gradually decreased as coil thickness increased from 0.1 mm to 0.5 mm. Distinctively, the mean kink radius from a coil pitch of 3 mm to 2 mm decreased from 9.2 ± 0.8 mm to 7.1 ± 0.7 mm, and at a coil pitch of 1 mm, the mean kink radius increased to 8.1 ± 0.9 mm. Sensitivity analysis revealed that kink radii were most dependent on coil pitch (Table 2). Across the bent buckling models, the average solve time was 05:35:33 ± 01:58:54 (days:hours:minutes), and the longest model completed in ∼10 days.

### 3.3 Candidate Graft Identification via Device Design Optimization

Results from the optimization function (Eq. 3) highlighted that unique graft designs could be identified across the graft parameter design space based on defined weighting values for the competing mechanical metrics. To assess that the function faithfully weighed the competing metrics, optimized design parameter combinations were identified for each singular metric. For example, based on a weighting of *α*= 1 and *β*= 0, the optimization function identified the coil-reinforced graft with design parameters of *δ* = 0.1 mm (thinnest coil thickness), *μ*= Bionate® 80A (softest coil material), and *p*= 10 mm (largest coil pitch, Figure 6), which corresponded to the most distensible graft model evaluated (Figure 5a). Similarly, a weighting of unity for the buckling load and kink radius terms confirmed design parameters that corresponded to the highest buckling load and smallest kink radius, respectively (Figure 6 and Figure. 5b, 5c).

**Figure 6.**
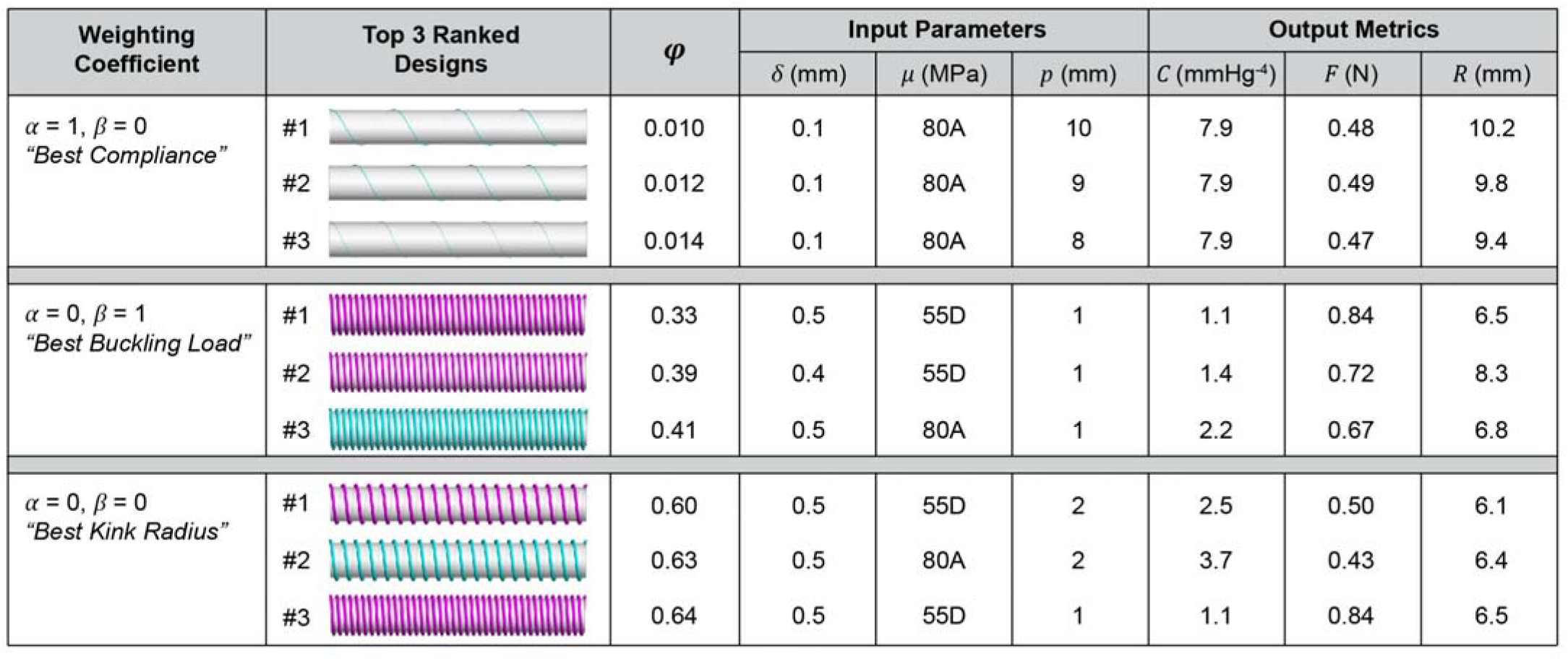
Candidate graft designs based on the optimization of a single mechanical behavior. Each block summarizes the weighting coefficient, illustrations of the top three ranked graft designs, corresponding optimization value (*φ*), input design parameters, predicted compliance (*C*), buckling load (*F*), and kink radius (*R*). [double-column]

Utilizing mixed weighting coefficients (i.e., evaluation of 2 or 3 metrics) led to unique coil-reinforced graft designs that integrated observations as when single metrics were weighted. For example, the top 2 optimized grafts when compliance and buckling load were equally weighted (i.e., *α*= *β* = 0.5) had the smallest coil thickness (0.1 mm) and softest coil modulus (Bionate® 80A) but different coil pitches (see Supplementary Figure 1). Only weighting the buckling load and kink radius (i.e., *α*= 0, *β*= 0.5) resulted in optimized grafts that had a coil thickness of 0.5 and a tight coil spacing (1 or 2 mm). Similar trends were observed when compliance and kink radius were weighted at 0.5 (i.e., *α*= 0.5, *β*= 0). Equal weighting (i.e., *α*= *β*= 0.33, Figure 7) or mixed weighting (e.g., *α*= 0.5, *β*= 0.25, see Supplementary Figure 2) across the mechanical metrics resulted in noticeable trends. For example, increasing the weighting coefficient on compliance resulted in grafts with thinner coils, comprised of a softer material, with a larger spacing. In contrast, increasing the weighting on the buckling resistance led to optimized graft designs with thicker coils and tighter spacing. Importantly, grafts with similar optimization function values can be achieved with differing graft designs and output metrics. With equal weighting across the mechanical metrics, for example, the 1^st^ and 3^rd^ ranked optimized graft designs had function values of 0.56 and 0.57, respectively, and coil spacings of 9 and 1 mm, respectively (Figure 8).

**Figure 7.**
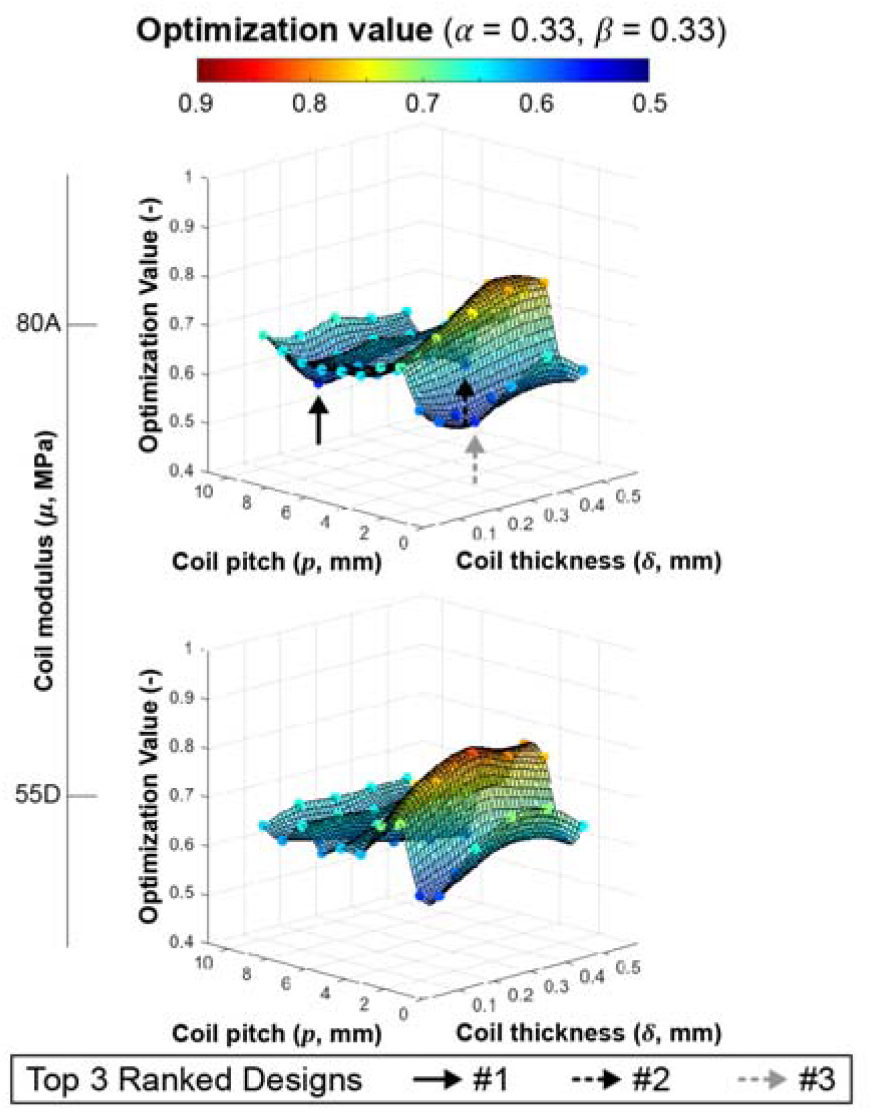
Design space evaluations for the 100 unique coil-reinforced graft combinations. Data were interpolated within the design space to evaluate the optimization function with equal weighting coefficients (*α* = *β*= 0.33). The optimization value was plotted against the – coil thickness (*δ*) and coil pitch (*p*). The top and bottom rows separately represent the two coil moduli (*μ*), Bionate ® 55D and 80A, respectively. The optimization function locates the global minimum as identified by the arrows representing the top 3 ranked designs for each output metric. [single-column]

**Figure 8.**
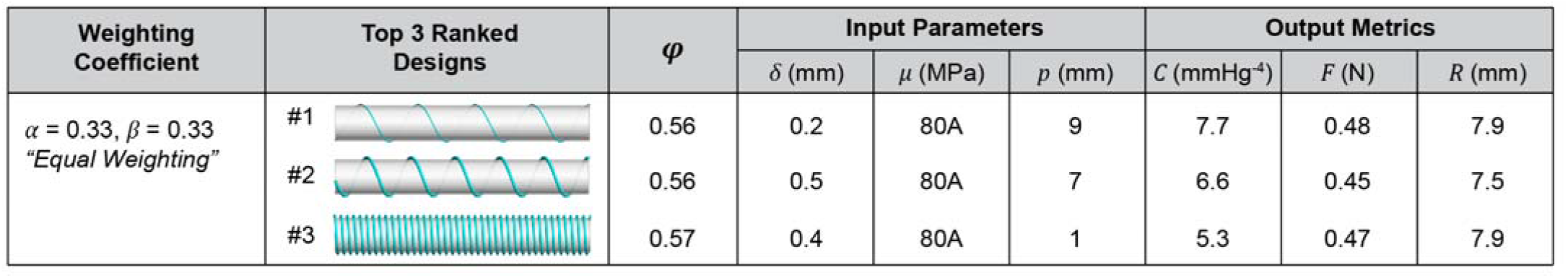
Candidate graft designs based on the optimization of three mechanical behaviors. Each block summarizes the weighting coefficient, illustrations of the top three ranked graft designs, corresponding optimization value (*φ*), input design parameters, predicted compliance (*C*), buckling load (*F*), and kink radius (*R*). [double-column]

## 4. DISCUSSION

This study presents a computational framework to optimize the mechanical behavior of synthetic vascular grafts. The framework parameterized the graft design (inputs), and FE models predicted graft compliance, buckling load, and kink radius (outputs) across the parameter space. Model predictions demonstrated agreement with experimental data, providing FE model validation across distinct and physiologically relevant loading conditions. Integrating the mechanical metrics into an optimization function reconciled competing biomechanical properties and efficiently identified candidate graft designs. Importantly, evidence of model credibility supports the use of modeling predictions and the presented framework, promoting model-directed design to accelerate graft optimization.

Computational modeling provides an *in silico* testbed to conduct studies that are time-consuming, challenging, or expensive to perform experimentally. In the presented study, we leveraged FE modeling to accelerate vascular graft design by systematically evaluating design input parameters and resulting mechanical response. Highlighting the merit of modeling in graft design, computational analysis of 100 parameter combinations utilizing HPC took 10 days to simulate with a cumulative time of 788:10:17 (days:hours:minutes). In contrast, experimental fabrication and benchtop testing of all graft combinations would have taken 6-9 months. This model-directed design approach utilized similar strategies developed and applied to optimize medical devices for vascular applications, including stents (Timmins, et al., 2007), coupling devices (Li, Agarwal, Coats, & Gale, 2016), and tissue-engineered graft design (Furdella, et al., 2020; Szafron, Ramachandra, Breuer, Marsden, & Humprey, 2019; Marsden, 2014). Important to the presented framework is recognizing the breadth of experimental approaches for graft fabrication and their impact on graft mechanical behavior. For example, modulating electrospinning techniques allows the manufacturing of unique corrugation patterns with varied compliance values (Akbari, Mohebbi-Kalhori, & Samimi, 2020). Custom 3D printed techniques allow the inclusion of distinct spiral reinforcement patterns (e.g., cross-pattern, ring, and spiral design) within a graft construct, and each pattern modifies the anti-kinking behavior of the graft (Wu D. J., et al., 2020). Towards advancing model-directed design, computational frameworks must be sufficiently modular and scalable to integrate novel fabrication approaches and coil designs, as well as mechanical behaviors (e.g., burst pressure, suture retention). Thus, the continued advancement of computational strategies is needed to realize their full potential (Cosgriff-Hernandez & Timmins, 2022).

The presented computational framework holds great potential in elucidating the complex mechanical determinants of graft performance that would be difficult and cost prohibitive to conduct experimentally. The computational models to simulate and analyze pressurization (compliance), axial buckling (buckling load), and bent buckling (kink radius) provided unique insight into the structure-function relationships of the complex structure. Researchers have previously performed extensive graft fabrication and testing (Bode, Mueller, Zernetsch, & Glasmacher, 2015). However, due to the inherent variability in sample fabrication and testing, as well as systematic and random error in experimental measurements, interpreting differences across graft designs is difficult, if not impossible. This experimental variability often confounds or limits attempts to establish structure-property relationships to guide graft design. In contrast, the presented computational predictions and sensitivity analysis readily informed differences across the graft design parameter space (See Figure 5, Table 2). For example, model predictions showed that the buckling load and kink radius were highly sensitive to coil pitch, whereas coil modulus had the least impact across all mechanical metrics. Therefore, fabrication strategies could be modified *a priori* to ensure fabricated grafts achieve the desired mechanical outcomes, thus saving considerable time and effort.

Experimental validation is essential for the applicability of any computational framework. The validation process aims to assess the predictive capability of the model for its intended use – a subtle point that needs emphasis (ASME Committee, 2006). The motivation for developing models is to make predictions for model applications where no experimental data could, or would, be obtained. This study presented a set of well-designed experimental tests that faithfully reproduced the underlying physics of the model and quantified the metrics of interest. Importantly, the chosen validation metrics were determined explicitly from the FE analysis and experiments, and each has a direct implication on graft regulatory approval and biological function. In addition, the low intra-sample errors in the experimental data support the choice of validation metrics, demonstrating repeatability and reliability (See Figure 4) (ASME Committee, 2006). Although studies have integrated FE modeling into the design of coil-reinforced grafts and reported graft deformations, stresses, and strains under lumen pressurization (Adhikari, Zimmerman, Dimble, Tucker, & Thomas, 2021), validating such quantities experimentally presents challenges due to measurement accuracy and resolution. Moreover, optimizing parameters based on FE-derived stress–strain values is difficult (Ovcharenko, et al., 2019). Hence, lower-order data (e.g., global displacements, clamp-to-clamp strains), which may be easier to obtain, are integral to model validation and establishing a higher level of model credibility (Anderson, Ellis, & Weiss, 2007). Collectively, model verification and validation, as well as uncertainty quantification (i.e., VVUQ), are paramount to establishing model credibility.

Although this computational framework has shown promise in the model-directed design of vascular grafts, there are limitations in the presented work that should be acknowledged. First, we assumed that the hydrogel-coated electrospun polyurethane graft composite behaved as an isotropic material (Eq. 1), which likely contributed to the differences between the FE and experimental data for buckling load and kink radius (See Figure 4b, 4c). The presented study largely focused on developing a unique computational framework for graft design, which provides the foundation to develop more advanced material models that account for the hydrogel phase and mesh phase (anisotropic fibers) into a ‘meso’-structural constitutive model that could be integrated into the pipeline (Motiwale, et al., 2022). Second, the FE models to evaluate compliance and buckling-resistance were evaluated as three individual models with idealized loading conditions. Although this approach afforded the ability to identify relevant trends based on changes in graft design, more complex loading conditions that simulate, for example, combined bend-twist buckling or helical buckling could offer additional insight into the graft response to *in vivo* loads. Lastly, the sensitivity analysis conducted herein (Eq. 4, Table 3) was performed with no additional computational expense and dictated by constraints. Future investigations that couple modern and certifiable uncertainty quantification techniques with FE approaches will provide a more rigorous approach to characterize model output sensitivities (Berggren, et al., 2024).

## 5. CONCLUSION

In conclusion, we present a computational approach to evaluate and optimize the mechanical behavior of coil-reinforced vascular grafts under physiologic loading conditions. Through complementary experiments, we demonstrate the accuracy of the FE models and the importance of evaluating distinct mechanical metrics relevant to graft success. Coupling the FE models with batch-processing and optimization schemes afforded the ability to efficiently evaluate 100 graft designs and reconcile three competing solid mechanical concerns. Future studies applying the presented model-directed design approach have the potential to promote the performance and success of synthetic vascular grafts without the need for time-consuming and costly experimental strategies.

## Abbreviations

CABG: Coronary Artery Bypass Grafting
NH: Neointimal hyperplasia
ePTFE: expanded Polytetrafluoroethylene
FE: finite element
NAGA: N-acryloyl glycinamide
HPC: High performance computing
VVUQ: Verification, validation, and uncertainty quantification

## CRediT author contribution statement

**David Jiang:** Methodology, Validation, Formal analysis, Writing - Original Draft, Review, & Editing, Visualization; **Andrew J. Robinson:** Methodology, Validation, Formal analysis, Writing – Review & Editing; Abbey Nkansah: Methodology; **Jonathan Leung:** Resources; **Leopold Guo:** Resources; Steve A. Maas: Software, Writing – Review & Editing; **Jeffrey A. Weiss:** Software, Writing – Review & Editing; **Elizabeth M. Cosgriff-Hernandez:** Conceptualization, Formal analysis, Writing – Review & Editing, Supervision, Project administration; **Lucas H. Timmins:** Conceptualization, Methodology, Formal analysis, Writing - Original Draft, Supervision, Project administration, Funding acquisition

## Acknowledgements

This work was supported, in part, by the National Institutes of Health (L.H. Timmins: R01 HL150608, J.A. Weiss: R01 GM083925, U24 EB029007). The authors thank DSM Biomedical for providing polyurethane samples for mechanical testing. The authors thank the Scientific Computing Institute at the University of Utah for access to their high-performance computational systems, which were supported by the National Institutes of Health (P41 GM103545, R24 GM136986).

## APPENDIX A. Mesh Convergence Study

A mesh convergence study was performed by refining the mesh size of the graft domain in the circumferential, axial, and radial directions (Fig. A.1a). Specifically, the number of circumferential elements(*n*_*θ*_) ranged from 8 to 64 elements (8 equally-spaced values). Next, the number of axial elements(*n*_*z*_) was determined by maintaining a 1:1 aspect ratio - the circumferential length of the element was equal to the axial length. Lastly, the number of radial elements (*n*_*z*_) were 1, 2, or 3 element(s) thick. Each model was analyzed to evaluate compliance, buckling load, and kink radius (Fig. A.1b-d). Based on the criteria of <3% change in each evaluated metric, results showed that a graft mesh domain of *n*_*θ*_ = 64, *n*_*z*_= 147,*n*_r_= 2(18,816 hexahedral elements) was sufficient to achieve convergence. The mean graft element volume was approximately 8.75 × 10^−3^ mm^3^.

**Figure A1.**
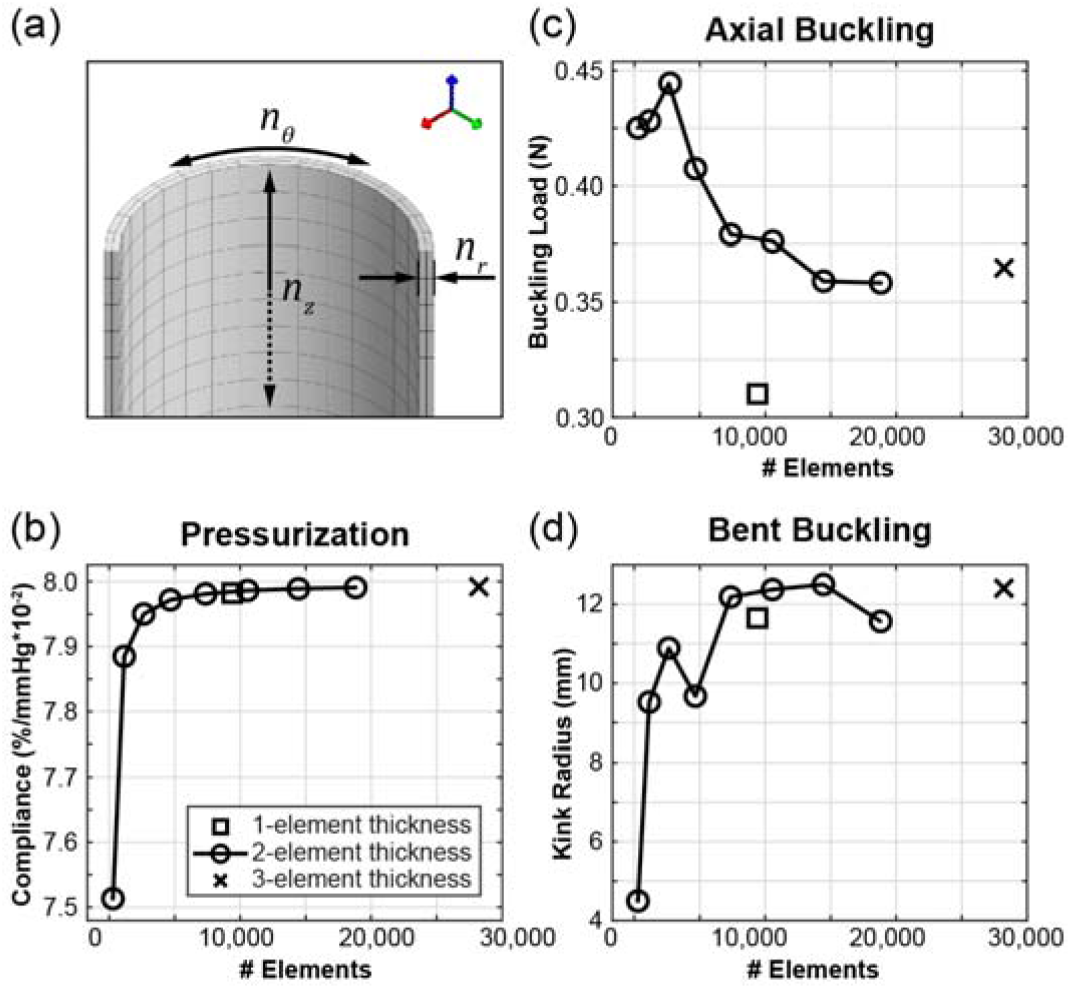
Mesh convergence study. (a) Variations in number of hexahedral elements composing the graft domain in the circumferential, axial, and radial directions. A convergence criterion of <3% change in evaluated metrics for (b) compliance, (c) buckling load, and (d) kink radius. [single-column]

## APPENDIX B. Energy Balance in Explicit Quasi-Static Analysis

The appropriate quasi-static response for the simulations was evaluated using the energy balance equation:

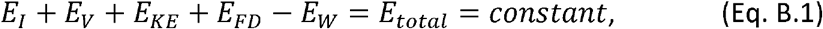

where *E*_1_ is the internal elastic strain-energy, *E*_*V*_ is the energy absorbed by viscous dissipation, *E*_*KE*_ is the kinetic energy, *E*_*FD*_ is the energy absorbed by frictional dissipation, *E*_*W*_ is the work of external forces, and *E*_*total*_ is the total energy in the system. To determine if the simulation was quasi-static (i.e., inertial forces were negligible), the work applied by the external forces was nearly equal to the internal energy of the system. The viscously dissipated energy was generally small due to the material damping. The kinetic energy of the deforming coil-reinforced graft was small – it did not exceed 5% of its internal energy throughout most of the analysis. The energy history for the axial buckling and bent buckling analyses are shown in Figure B.1.

**Figure B1.**
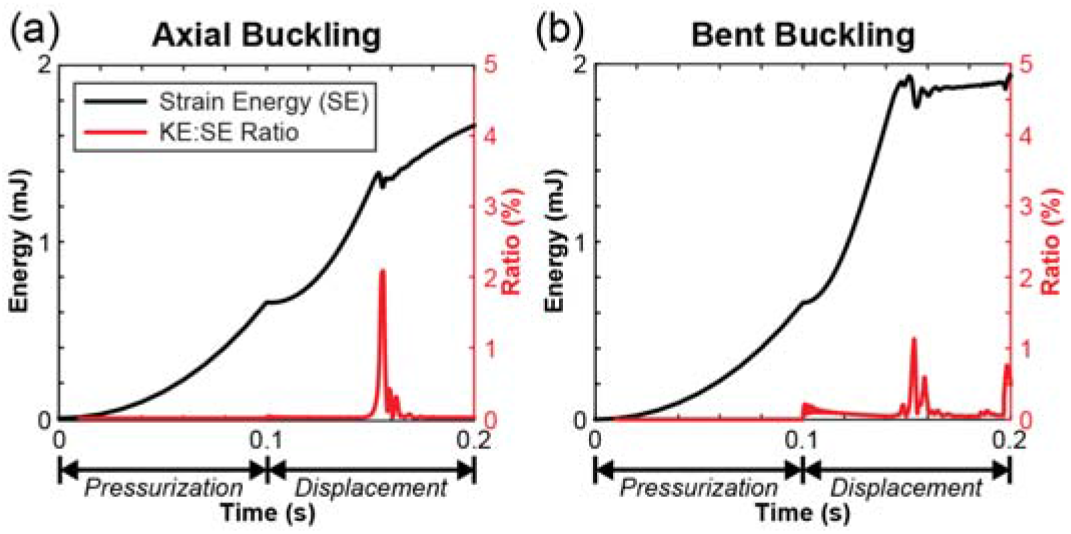
Energy history for the explicit quasi-static analysis. (a) Axial buckling energy history. (b) Bent buckling energy history. The ratio of kinetic energy to internal strain energy did not exceed 5% throughout the analysis. A minor increase in kinetic energy occurred during the buckling event. [single-column]

## Footnotes

2 www.febio.org

3 www.gibboncode.org

4 See Appendix A

5 The explicit time-integration scheme in `FEBio` utilizes a “lumped” mass matrix, where the matrix elements of each row are summed together and assigned to the diagonal.

6 See Appendix B

