## Supplementary Data for "A Computational Framework to Optimize the Mechanical Behavior of Synthetic Vascular Grafts"

### **Supplementary Materials**

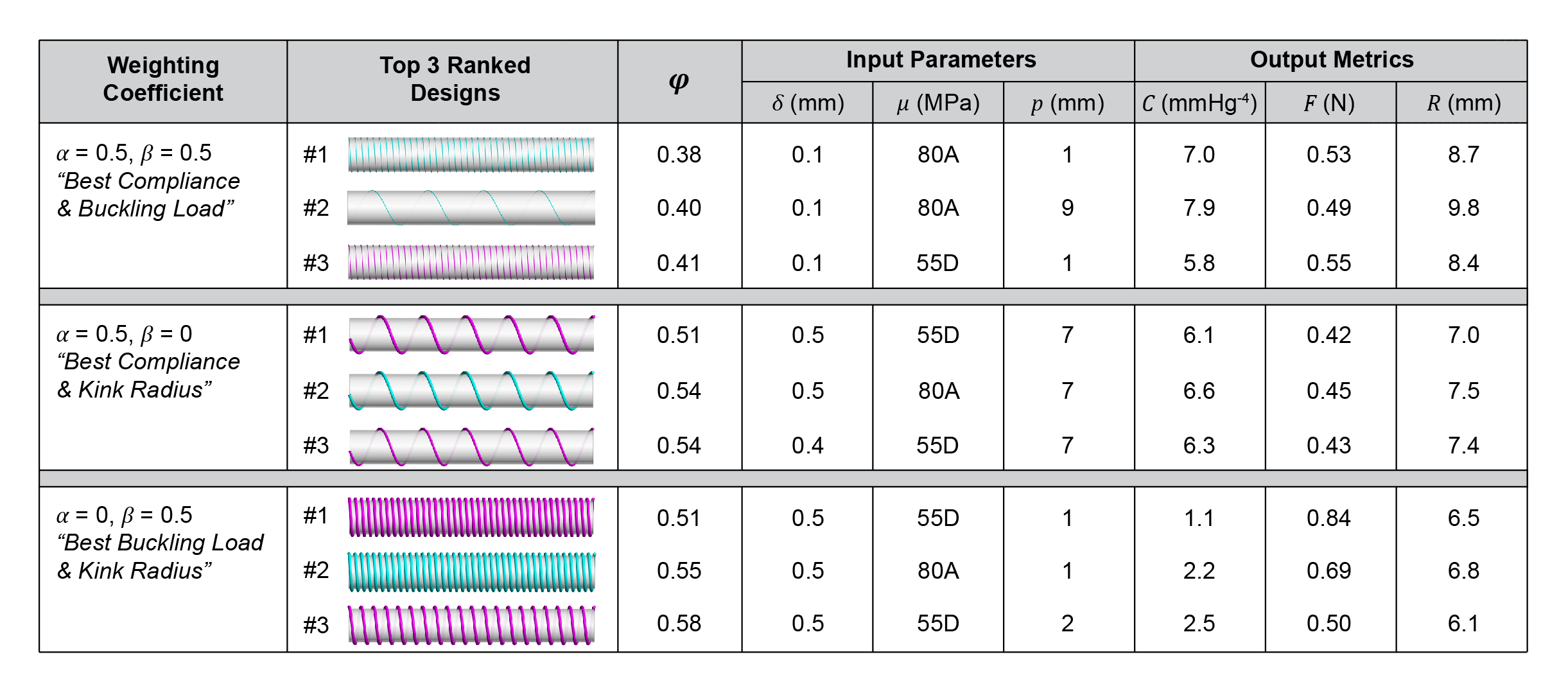

**Supplementary Figure 1.** Candidate graft designs based on the optimization of two mechanical behaviors. Each block summarizes the weighting coefficient, illustrations of the top three ranked graft designs, corresponding optimization value ($\varphi$), input design parameters, predicted compliance ($C$), buckling load ($F$), and kink radius ($R$).

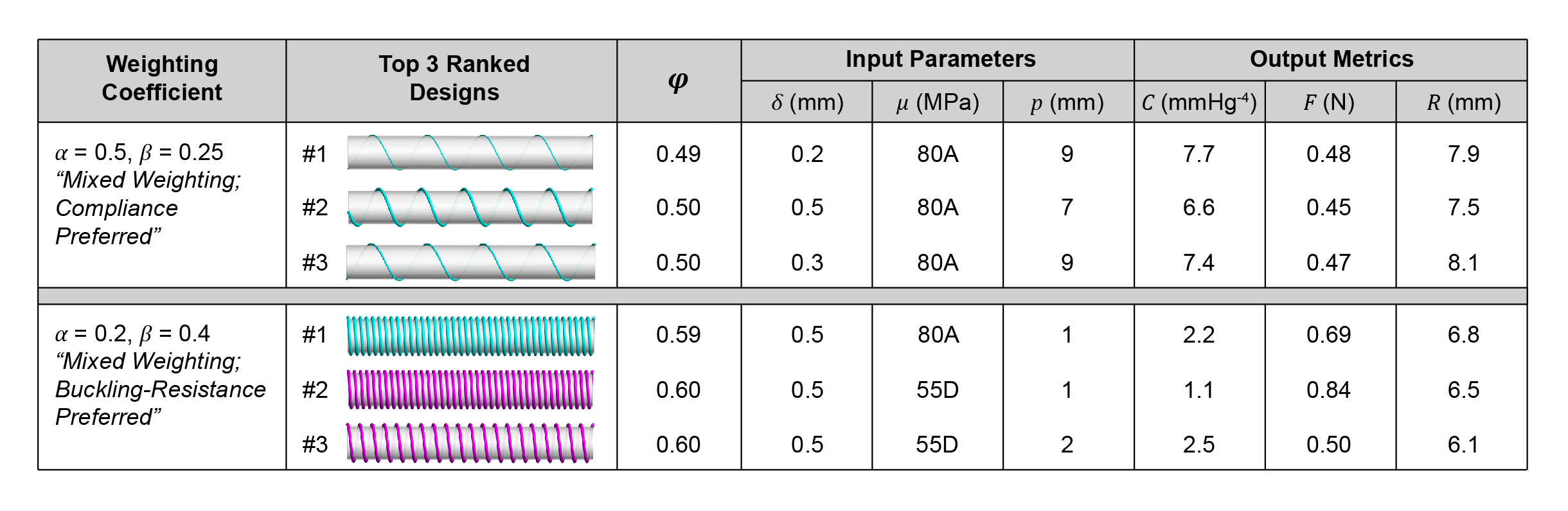

**Supplementary Figure 2.** Candidate graft designs based on the optimization of three mechanical behaviors. Each block summarizes the weighting coefficient, illustrations of the top three ranked graft designs, corresponding optimization value ($\varphi$), input design parameters, predicted compliance ($C$), buckling load ($F$), and kink radius ($R$).
